# The genome-wide mutational consequences of DNA hypomethylation

**DOI:** 10.1101/2022.10.04.510776

**Authors:** Nicolle Besselink, Janneke Keijer, Carlo Vermeulen, Sander Boymans, Jeroen de Ridder, Arne van Hoeck, Edwin Cuppen, Ewart Kuijk

## Abstract

DNA methylation is important for establishing and maintaining cell identity and for genomic stability. This is achieved by regulating the accessibility of regulatory and transcriptional elements and the compaction of subtelomeric, centromeric, and other inactive genomic regions. Carcinogenesis is accompanied by a global loss in DNA methylation, which facilitates the transformation of cells. Cancer hypomethylation may also cause genomic instability, for example through interference with the protective function of telomeres and centromeres. However, understanding the role(s) of hypomethylation in tumor evolution is incomplete because the precise mutational consequences of global hypomethylation have thus far not been systematically assessed. Here we made genome-wide inventories of all possible genetic variation that accumulates in single cells upon the long-term global hypomethylation by CRISPR/CAS9-mediated conditional knockdown of *DNMT1*. Depletion of DNMT1 results in a genomewide reduction in DNA methylation levels. Hypomethylated cells show reduced proliferation rates, reactivation of the inactive X-chromosome and abnormal nuclear morphologies. Prolonged hypomethylation is accompanied by increased chromosomal instability. However, there is no increase in mutational burden, enrichment for certain mutational signatures or structural changes to the genome. We conclude that the primary consequence of global hypomethylation is chromosomal instability and does not necessarily lead to other small-scale or structural mutational effects.

## Introduction

Epigenetic mechanisms confer cell identity by regulating gene activity, allowing cells to have different phenotypes while sharing the same genotype[1]. To switch phenotypes cells need to remodel their epigenetic landscapes, for example during differentiation[2], reprogramming[3], or the transformation of healthy cells into tumor cells[4]. A key epigenetic modification in mammalian cells is DNA methylation, the covalent attachment of a methyl group to the 5th carbon atom of cytosine. In healthy human cells, around 85% of the cytosines that are flanked by guanines (CpG sites) are methylated. Notable exceptions are CpG islands (CGIs), regions with high GC content that are mostly unmethylated (<10%) and are associated with active promoter regions of housekeeping genes and tumor suppressor genes (TSGs) [5].

The DNA methylation landscape is sculpted by the joint activity of DNA methyltransferases (DNMT) and ten-eleven translocation (TET) enzymes. DNMT3A and DNMT3B are responsible for the de novo establishment of DNA methylation. DNMT1 maintains DNA methylation in dividing cells and is supported by UHRF1 that recognizes hemi-methylated substrates at replication forks[6]. Active demethylation is accomplished by TET1, TET2, and TET3, through the hydroxylation of methylcytosine, followed by oxidation of hydroxymethylcytosine to 5-formylcytosine (5fC) and then to 5-carboxycytosine (5caC). These bases are subsequently recognized by base excision repair and replaced by cytosine. DNMT and TET enzymes are frequently amplified or mutated in cancer and can have a causal role in carcinogenesis [7–14].

Many studies have provided evidence that genetically or chemically induced loss of DNA methylation results in genomic instability [15–20]. DNA methylation supports genomic stability by promoting DNA condensation and the inhibition of transcription of constitutive heterochromatic genomic elements including centromeric and pericentric heterochromatin, LINE elements and subtelomeric regions [21,22]. Tumorigenesis is generally accompanied by a global decrease in DNA methylation [23,24], which leads to the concomitant deprotection of these heterochromatic regions. Pericentric heterochromatin, LINE elements and subtelomeric regions are frequently affected in cancer [25–28], indicating that the cancer-associated changes to the DNA methylation landscape coincide with genome instability. Thus, the epigenetic instability of transforming cells may interfere with the protective role of DNA methylation in maintaining genomic integrity, leading to genomic instability. However, the genetic - epigenetic relationship is complex and our knowledge on the genomic consequences of the epigenetic reprogramming that coincide with cancer development is far from complete.

To improve our view on cancer evolution and tumor heterogeneity we need a better understanding of how reprogramming of the epigenetic software influences the genetic hardware. In recent years, whole genome sequencing of cancer genomes has increasingly been used for the identification of mutational signatures, distinct patterns of mutation accumulation that provide insight into past mutational processes, such as DNA repair deficiencies, endogenous mutational processes, and exposure to exogenous mutagens such as tobacco-smoking, UV-light or anti-cancer treatments [29]. Over 100 mutational signatures have been described that involve single, double and clustered base substitution signatures and indel signatures[30,31]. In addition, sixteen signatures have been identified that are based on patterns of structural variation such as deletions, tandem duplications, and inversions [32]. The etiology for many mutational signatures is still unknown.

In spite of the myriad genomic consequences that may follow DNA methylation loss, the genomic consequences of long-term DNA hypomethylation have thus far not been systematically characterized and no mutational signatures have been linked to DNA methylation loss. In this study, we aimed to fully characterize all genomic consequences of global methylation loss; from the whole chromosome level down to single base resolution. We performed CRISPR/CAS9 mediated conditional knockdowns of *DNMT1* and examined the genetic consequences after prolonged culture by whole genome sequencing at 30X coverage of expanded clones to capture events at high resolution affecting few or single bases and by single cell DNA sequencing to capture chromosome scale events.

## Results

For these experiments we made use of a human female TERT immortalized RPE-1 cell line, which lacks a transformed phenotype and is near diploid. Because of its key role in maintenance of methylation, interference with DNMT1 activity will lead to a progressive loss of DNA methylation in cycling cells. In most cell lines, *DNMT1* is an essential gene [33,34]. In line with these observations, our attempts to create a *DNMT1* knockout in RPE-1 cells failed, while knockouts for non-essential genes were successful (data not shown). Therefore, we decided to use CRISPR interference (CRISPRi) to knock down *DNMT1* gene expression[35]. In CRISPRi, a nuclease dead version of Cas9 (dCAS9) fused to the transcriptional inhibitor KRAB[36] (dCAS9-KRAB) is guided to the transcriptional start site by a sgRNA to block the transcription machinery resulting in decreased expression of the gene of interest. In combination with a doxycycline inducible sgRNA, CRISPRi allows temporal downregulation in full isogenic cell lines.

We established a clonal hTERT RPE-1 P53^-/-^ dCAS9-KRAB doxycycline inducible sgRNA^*DNMT1*^ cell line (hereafter referred to as *DNMT1kd* cells). A *P53*^*-/-*^ line was used to increase the permissiveness of cells for all types of mutations and avoid loss of such events due to P53-induced senescence or apoptosis. *DNMT1kd* cells showed clear upregulation of the sgRNA upon addition of doxycycline, which was followed by long-term downregulation of *DNMT1* at the mRNA and protein levels (Fig. 1a-c). Methylation arrays covering over 850,000 methylation sites revealed a genome wide reduction in methylation levels in *DNMT1* knockdown cells in three independent experiments (Fig.1d). The loss was equally distributed over genes, promoters and CpG islands. Prolonged downregulation of *DNMT1* delayed growth by approximately 50%, without leading to a full cell cycle arrest (Fig. 2a), which is in line with previous observations [19,20]. Further characterization of *DNMT1kd* cells by RNA seq confirmed robust knockdown of *DNMT1* and further revealed that global hypomethylation resulted in a relative increase in gene activity (Fig. 2b). In particular, genes on the X-chromosome were upregulated, indicating partial reactivation of the inactive X-chromosome (Fig. 2c). Moreover, doxycycline treated cells acquired an abnormal kidney-shaped nuclear morphology, indicating that DNA methylation loss disturbs global genome organization (Fig. 2d). CRISPRi mediated knockdown of *DNMT1* in RPE-1 cells is a relatively clean method to reduce DNA methylation compared to alternative methods such as short hairpin mediated knockdown or chemical inhibition that have more risk of adverse side-effects unrelated to the loss of DNA methylation and are generally restricted to short-term consequences[37–39]. Together, this conditional *DNMT1kd* cell line is a powerful tool to study the long-term consequences of DNA methylation loss.

**Figure 1:**
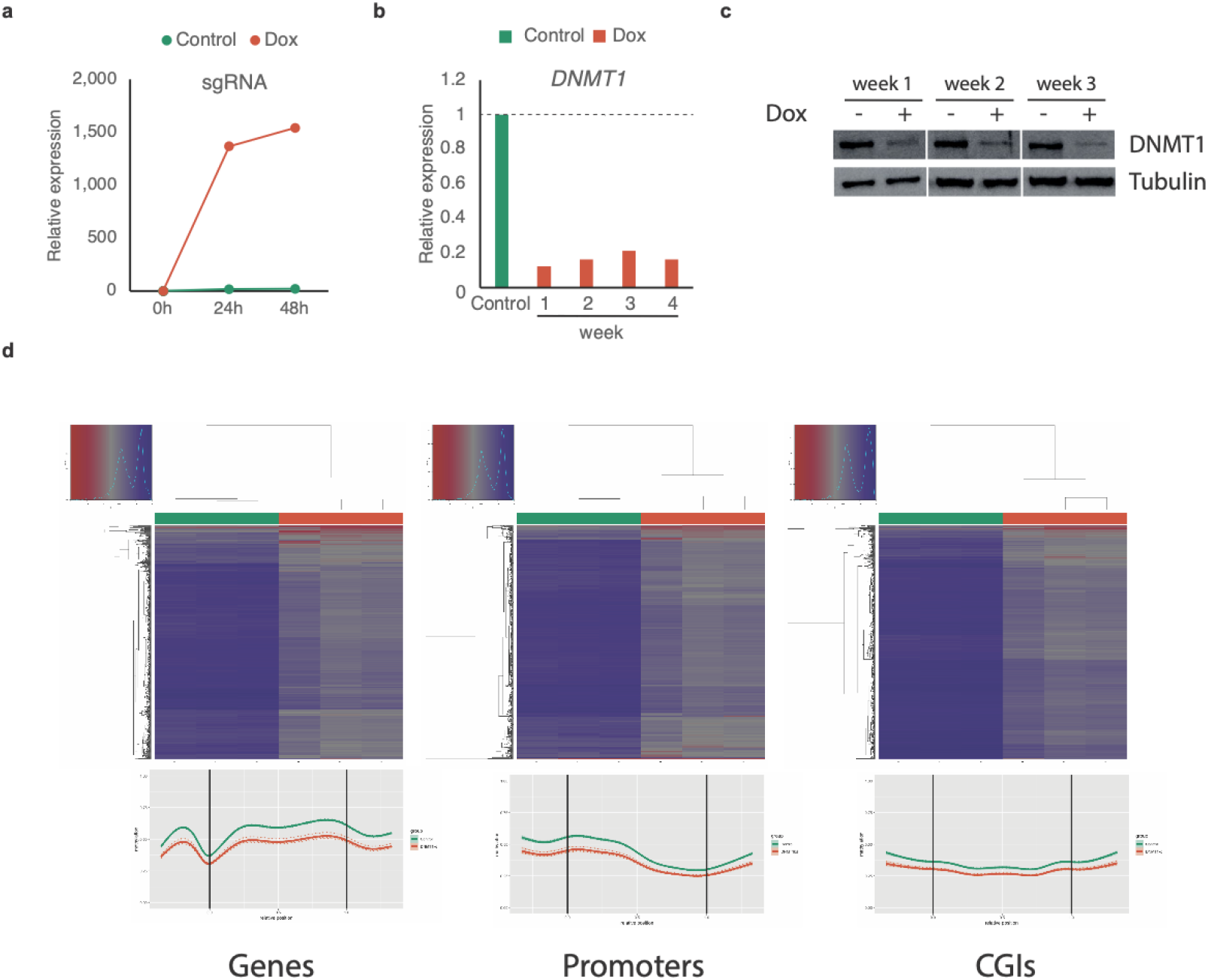
Characterization of *DNMT1kd* cells. (a) Relative increase in doxycycline-inducible sgRNA expression upon 24h and 48h hours of treatment with doxycycline.(b) Relative *DNMT1* expression levels upon prolonged doxycycline treatment of *DNMT1kd* cells.(c) DNMT1 protein expression levels upon prolonged doxycycline treatment of *DNMT1kd* cells.(d) Top panels: Unsupervised hierarchical clustering of samples based on methylation values. Notice distinct clustering of untreated *DNMT1kd* cells (green) versus doxycycline treated (red) *DNMT1kd* cells. Lower panels: Regional methylation profiles (composite plots) according to sample groups. *DNMT1kd* cells consistently show reduced methylation density represented by lower beta values. For each region in the corresponding region type, relative coordinates of 0 and 1 correspond to the start and end coordinates of that region respectively. Coordinates smaller than 0 and greater than 1 denote flanking regions normalized by region length. Horizontal lines indicate region boundaries.

**Figure 2:**
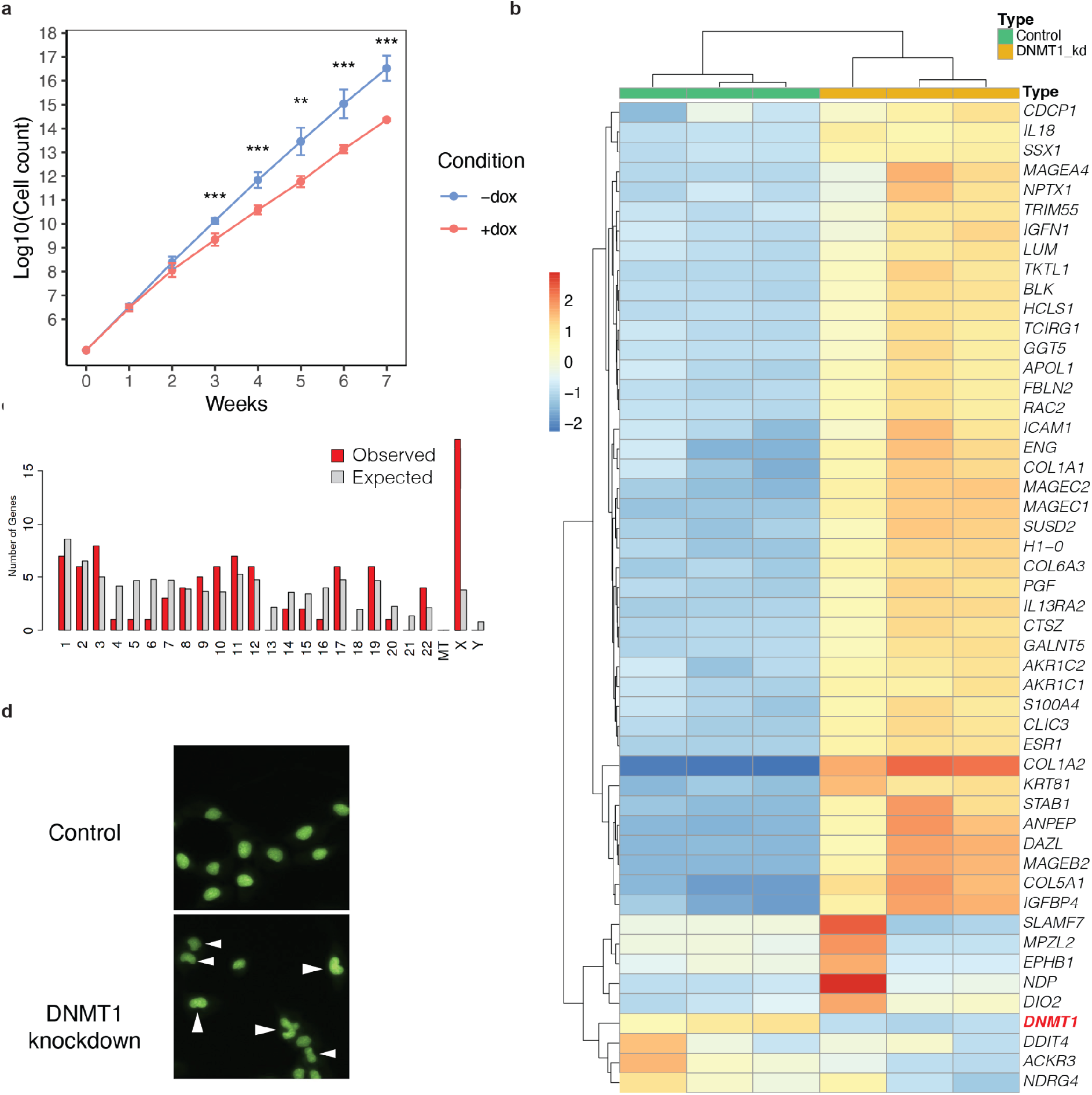
Phenotype of *DNMT1kd* cells. (a) Growth curves of doxycycline induced *DNMT1kd* cells and control cells (n=4 per condition per time-point). (b) Unsupervised hierarchical clustering heatmap of the 50 most differentially expressed genes between doxycycline induced *DNMT1kd* cells and control cells. (c) Distribution of genes on chromosomes as determined by ShinyGO 0.76[73]. Chi-square test p=5.8E-08. (d) Morphology of control cells and cells with hypomethylated genomes. Arrowheads denote cells with abnormal nuclear morphology.

To examine the genomic consequences of global DNA methylation loss in single cells at nucleotide resolution we performed prolonged downregulation of DNMT1 (6 weeks) to allow putative genetic changes to occur, followed by a clonal step. Whole genome sequencing at 30X coverage of expanded clones followed by computational analysis enables the identification of all types of genetic variation, including single base substitutions (SBSs) and indels that are private to each clone. We previously applied this highly sensitive method to identify and characterize the mutational signatures in individual cells in healthy, diseased, and perturbed conditions [40–45]. Loss of DNA methylation has previously been associated with mismatch repair (MMR) deficiency [46] and therefore we anticipated to find an elevated mutational burden as well as an increase in the contribution of MMR-related mutational signatures. In contrast to our expectations we did not observe an increase in SBS burden (Student’s t-test, *p* value=0.18; Fig. 3a) or in the number of indels (Student’s t-test, *p* value=0.33; Fig. 3b), although numbers tended to be lower in the hypomethylated clones when compared to the control condition, possibly reflecting the differences in proliferation rates (Fig. 2a). The mutational spectrum (Fig. 3c) and 96 mutational profiles (Fig. 3d-e) were highly similar between hypomethylated cells and control cells, indicating that reduced methylation did not lead to a shift in the activity of mutational processes. (Fig. 3f). Notably, in this study we relied on a sample size set (n = 3 per condition) similar to previous studies that also used manipulated tissue culture systems and which has been adequate for the detection of differences in mutation rates and to identify induced mutational patterns [40,43,47]. Structural variant (SVs) analysis revealed that most SVs were shared between clones of both conditions indicating these SVs were acquired prior to *DNMT1* knockdown. Notable exception was a chromothripsis event that affected the q-arm of Chromosome 19, which was only present in two of the three clones of the hypomethylated cells (Fig. 3g).

**Figure 3:**
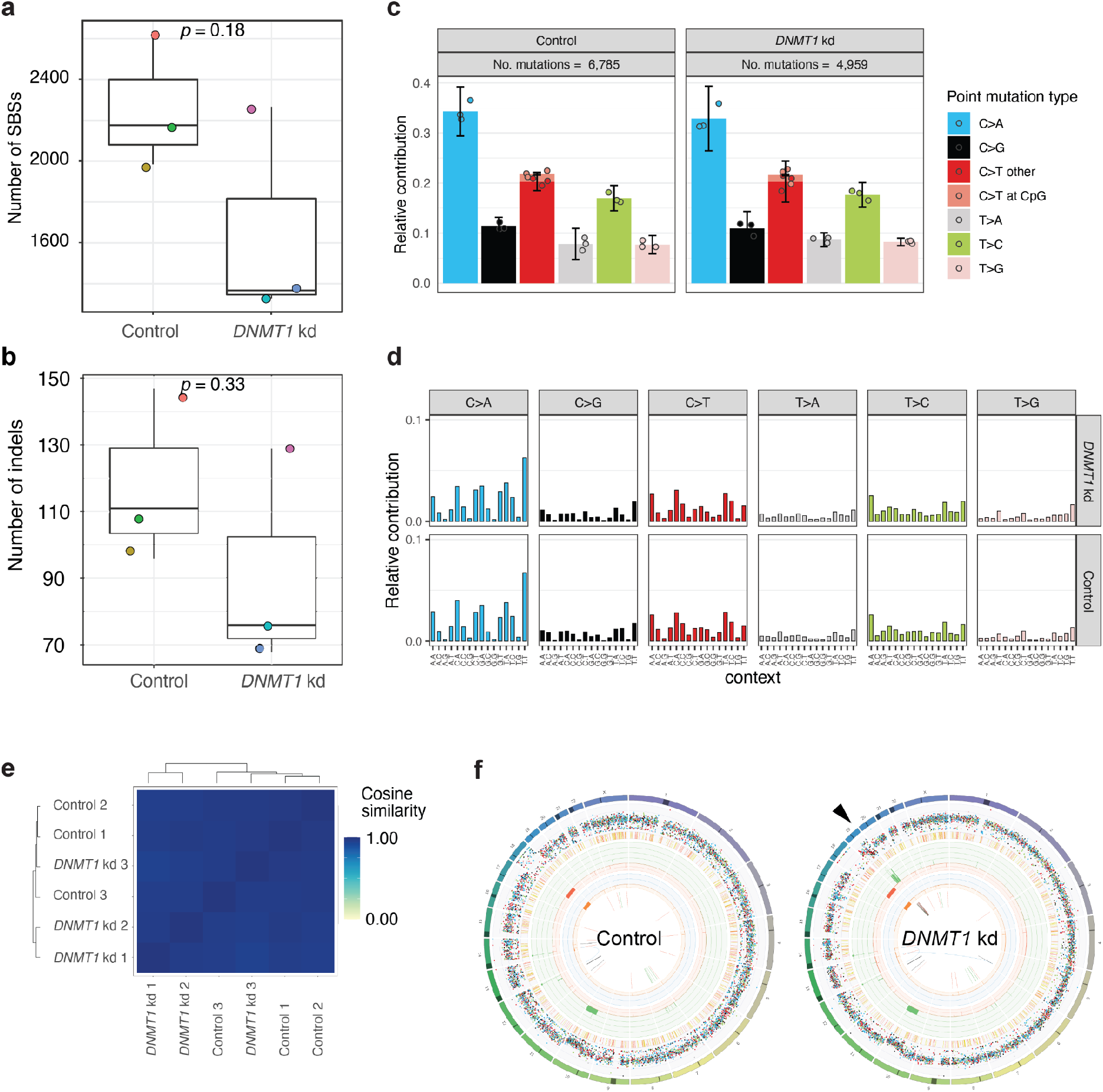
Mutational impact of global DNA methylation loss. (a) Box-plot with the individual data points of the number of single base substitutions per clone. Each clone is represented by a unique color. (b) Box-plot with the individual data points of the number of indels per clone. Each clone is represented by a unique color. Same color scheme as in a. (c) Average mutational spectra of (left) control clones and (right) hypomethylated DNMT1 knockdown clones. (d) Average mutational profiles of (top) hypomethylated DNMT1 knockdown clones and (bottom) control clones. (e) Cosine similarity between the mutational profiles of hypomethylated *DNMT1* knockdown clones and control cells. (f) Circos plots from a clone with (left) normal DNA methylation levels and (right) a hypomethylated *DNMT1* knockdown clone. Arrowhead denotes a chromothripsis event at the q-arm of chromosome 19, which was observed in 2 out of 3 clones.

Chromothripsis has mechanistically been linked to the missegregation of chromosomes or chromosome arms followed by damage and non-homologous repair in micronuclei[48]. The observed chromothripsis event may therefore be the result of increased chromosomal instability (CIN), as has been described previously for hypomethylated cells[49]. To further investigate CIN, we performed single cell DNA sequencing (Fig. 4a). Doxycycline treatment for 6 weeks of *DNMT1kd* cells resulted in increased aneuploidy and higher cellular heterogeneity (Fig. 4b). We reasoned that the increased chromosomal instability was the result of the loss of DNA methylation in peri-centromeric regions. However, the repetitive nature of these regions precludes assessment of DNA methylation by DNA methylation arrays. Therefore, we decided to perform long-read Nanopore sequencing of peri-centromeres, which allows both the direct measurement of methylated cytosines as well as unique mapping to the telomere-to-telomere reference genome[50]. Similar to the genomic regions that were assessed by methylation arrays, pericentromeric regions of doxycycline-treated cells showed a consistent reduction in DNA methylation levels (Fig. 4c).

**Figure 4:**
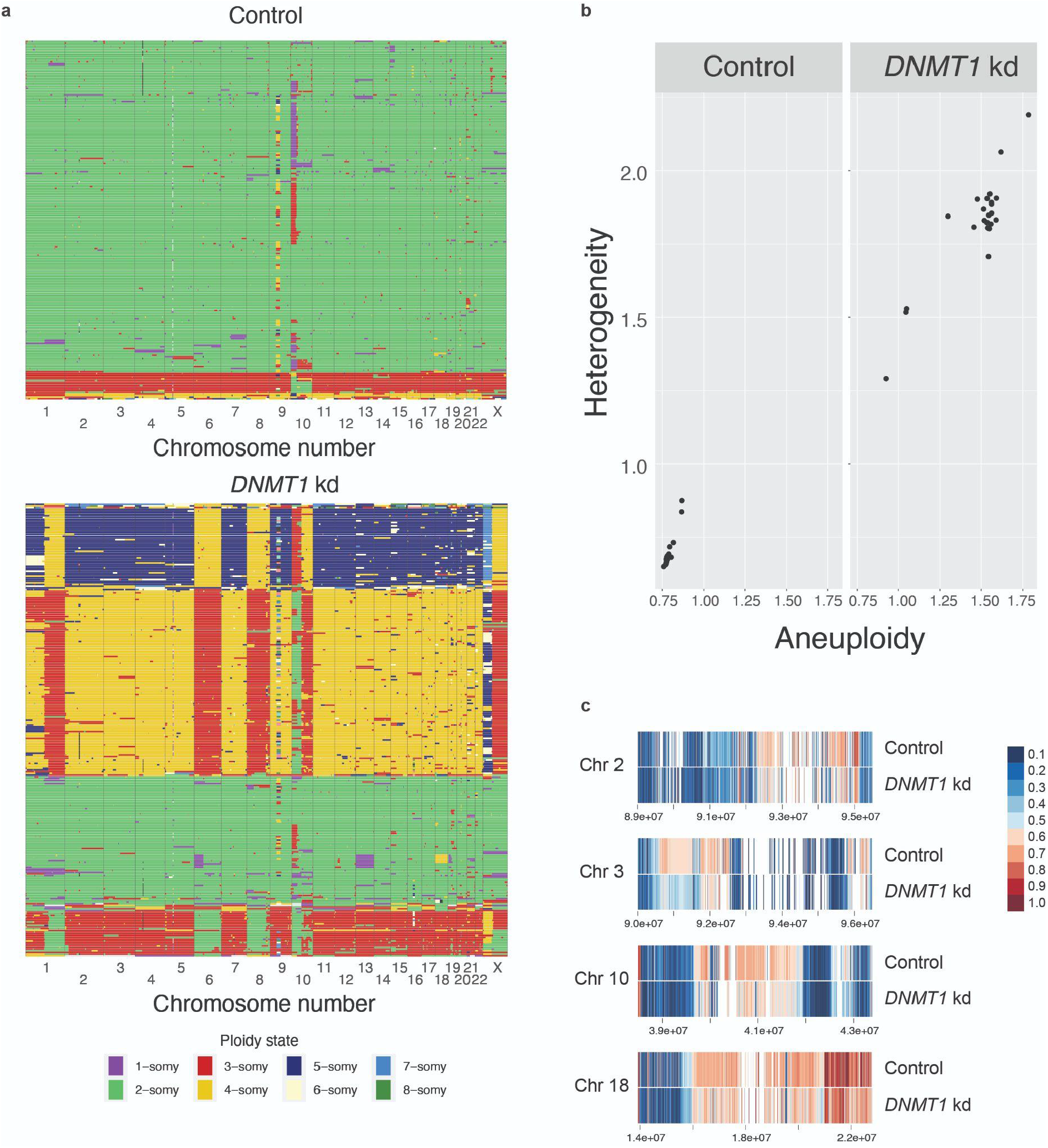
Impact of global DNA methylation loss on chromosomal instability. (a) Single cell DNA sequencing results showing the copy number state per chromosome per cell for (top) control clones and (bottom) hypomethylated DNMT1 knockdown clones. Each row represents a single cell and each column represents a chromosome. (b) Aneuploidy and heterogeneity scores for (left) control clones and (right) hypomethylated DNMT1 knockdown clones calculated with Aneufinder. Each dot represents a chromosome. (c) Reduced DNA methylation levels in pericentromeric regions of chromosomes 2, 3, 10, and 18 upon *DNMT1* knockdown as determined by Nanopore DNA sequencing. Coordinate values are indicated.

## Discussion and Conclusion

Ample evidence has established that DNA methylation is required for genomic integrity [15–20]. However, DNA methylation is associated with many different genomic elements providing different routes towards genomic instability upon methylation loss [51]. Reduced methylation can destabilize pericentromeric heterochromatin inducing rearrangements of chromosome arms, as has been described in for example Wilms tumors and hepatocellular carcinoma [52,53], and upon chemical interference of cell lines with the DNA hypomethylating drug 5-aza-2’-deoxycytidine [20]. Additionally, loss of methylation at repetitive elements may lead to genomic instability through secondary DNA structures that cause replication stress, DNA breaks and recombination between repeats [51]. Methylation loss can also lead to derepression of transposable elements that in turn can promote genomic rearrangements [26,54,55]. A decrease in methylation could also lead to more transcription and formation of R-loops, which are a source of replication stress, DNA breaks and genome instability [56]. Furthermore, reduced methylation at subtelomeric regions could lead to telomere instability [57]. Finally, DNA methylation has been implicated in DNA mismatch-repair [46] and the absence of DNA methylation may lead to mild MMR-deficiency.

Because of these possible effects of DNA methylation loss, we hypothesized that experimentally induced DNA demethylation would lead to a broad spectrum of genetic variation, ranging from SBSs, to indels, and from SVs to chromosomal aberrations. With a systematic approach, using highly sensitive methods to measure all types of genetic variation in individual cells [40,42,58], we identified only chromosomal instability as the foremost mutational consequence of DNA methylation loss.

There are several possible explanations for why we did not observe other types of genetic variation. In our model, DNA methylation loss was incomplete, so it is possible that more genetic events beyond chromosomal instability will occur if DNA methylation is further reduced, although this would likely severely impact on viability of the cells. Additionally, the hypomethylation effects may be cell-type specific. For example, the RPE-1 cell line that we used was immortalized by overexpression of *hTERT*, which may counteract any destabilization of telomeres caused by DNA methylation loss. Other cell types may be more sensitive to DNA hypomethylation in peri-telomeric regions. The effects of DNA methylation loss may also be context dependent and require additional circumstances. For example, global DNA demethylation may derepress transposons, but these may only become active when accompanied by the loss of repressive histone modifications and when post-transcriptional and post-translation defense mechanisms that suppress their mobility also fail. The mutational patterns induced by DNA methylation loss may also be too subtle to be detected above the background of more common mutational processes, e.g. those induced by culturing conditions [40]. Taken together, while our observations do not support an important role for DNA methylation in mutational processes other than CIN, subtle effects may be detected by capturing more variants, for example by extending the culture period of hypomethylated cells or by the analysis of vast amounts of clones expanded from single cells, which would be a very costly endeavor.

Hypomethylation induced CIN in cancer cells may contribute to tumor heterogeneity and cancer evolution. The exact mechanism through which hypomethylation causes CIN is as yet unknown. Proposed non-exclusive mechanisms include increased recombination of centromeric repeats, increased DNA breaks in centromeres, dysregulation of the centromeric protein network, increased transcription of α-satellite transcripts, defective assembly of centromeres/kinetochores and premature cohesion loss [22,32]. Future work may uncover the mechanistic link between pericentromeric hypomethylation and chromosomal instability.

## Materials and Methods

### Molecular cloning and virus production

We replaced the shRNA scaffold of pLV.FUTG.Tet.Inducible.shRNA with the sgRNA scaffold from lenticrispr v2[59], by InFusion cloning (Takara). Next, we annealed oligonucleotides for the guideRNA of *DNMT1* (Top: CACCGGGTACGCGCCGGCATCTCGG; Bottom: AAACCCGAGATGCCGGCGCGTACCC) and subsequently ligated the product into BsmbI digested pLV.TETi.sgRNA to obtain pLV.TETi.sgDNMT1. Selection of this sgRNA sequence was predicted to be highly effective for CRISPR interference based on chromatin, position, and sequence features [35]. To make RPE-1 cells more permissive to the accumulation of genetic variation, *TP53* knock-outs were generated with sgRNAs targeting the gene sequences 5’-GGGCAGCTACGGTTTCCGTC-3’. The annealed DNA oligonucleotides were cloned in px458 that also contains Cas9 followed by a 2A-EGFP sequence [60]. For virus production, lentiviral plasmids carrying the transgenes of interest were co-transfected with pRSV-REV, pMD2g and pMDLG-pRRE in HEK293T cells. Virus was harvested 3 days post-transfection and collected for immediate use or concentrated using Lenti-X Concentrator solution and stored at -80. Concentrated or undiluted virus was added to RPE-1 cells in a dilution series together with 4µg/ml polybrene. Successfully transduced cells were subsequently selected with the appropriate mammalian selection marker.

### Cell culture

hTERT RPE1 cells (ATCC) and HEK239T cells were cultured in DMEM, 10% fetal bovine serum, 1% penicillin, 1% streptomycin at 37°C, and 5% CO2. RPE1 *TP53*^*-/-*^ cells were transduced with lentivirus carrying lenti.EF1a.dCas9.KRAB.Puro [36] followed by transduction with pLV.TETi.sgDNMT1 lentivirus. RPE1-p53ko-dCas9-KRAB-Teti-DNMT1 cells were cultured with 10 ug/ml puromycin for selection of the dCas9-KRAB plasmid, 10 µg/ml blasticidin for selection of guide RNA plasmid. Clonal cell lines were established by limiting dilution series. The expression of the guideRNA was induced by stimulation with doxycyclin 2 µg/ml, which was refreshed every 2-3 days. Cell counts were performed every week and 5.3×10E+3 cells/cm^2^ were seeded. Cell counts were multiplied by split ratio to correct for passage. To establish hTERT-RPE1 CRISPR knock-out cell lines, cells were seeded in a 10 cm dish and transfected with vectors encoding both Cas9 and sgRNA target sequence using an Amaxa Nucleofector II instrument (Lonza). Forty-eight hours after transfection single GFP+ cells were sorted into 96-wells plates on FACS ARIA II/III Flow Cytometer (BD Biosciences) and expanded. Clones were tested for genome editing with PCR and Sanger sequencing and analyzed with the ICE CRISPR tool [61].

### RT-qPCR

RNA was isolated using the RNeasy Mini kit (Qiagen) and further purified using 3M NaAc and isopropanol if required. RT-qPCR was performed using Luna Universal One-Step RT-qPCR kit (NEB) using the following primers: *DNMT1* forward: 5’-AGCGGAGGTGTCCCAATATG-3’, *DNMT1* reverse: 5’-GAGACACAGTCCCCCACTTC-3’, sgRNA forward: TTTAGAGCTAGAAATAGC, sgRNA reverse: CGACTCGGTGCCACTTTTTC, *GAPDH* forward: 5’-AAATCCCATCACCATCTTCCAGGAGC-3’, *GAPDH* reverse: 5’-CATGGTTCACACCCATGACGAACA-3’. GAPDH was used as a reference gene and relative gene expression levels were calculated by ΔΔCt analysis[62], comparing cells cultured with doxycycline to cells cultured without doxycycline.

### Western Blot

Samples were collected in Laemmli buffer and incubated at 100°C for 10 min. Pageruler plus prestained protein ladder (ThermoFisher) and 20 ug of each sample were loaded on a 6% or 10% SDS-PAGE gel and transferred to a nitrocellulose membrane using Trans-Blot Turbo Transfer System (Bio-Rad). Western Blot was blocked using 5% ELK and incubated overnight with primary antibody mouse monoclonal anti-UHRF1 (Santa Cruz, SC-373750, 1:250) or rabbit polyclonal anti-DNMT1 (Invitrogen, PA3-16556, 1:1000) or monoclonal mouse anti-tubulin (Sigma-Aaldrich, T5168, 1:4000) and secondary HRP-conjugated antibody goat anti-mouse (1:2500) and goat anti-rabbit (1:2500). Blot was imaged using Amersham ECL (GE Healthcare) and Amersham Imager 600.

### Staining and microscopy

Cell culture medium was removed and 1 ml of staining solution, consisting of 2.5 uM SYTO 11 Green Fluorescent Nucleic Acid Stain (Invitrogen) in cell culture medium was added to each well of a 6-well plate and cells were incubated at 37°C for 60 min. Cells were imaged using an EVOS M5000 Imaging System at 10X magnification.

### Single cell DNA sequencing

Single cell sequencing was performed by the Single Cell Sequencing Core facility at the Hubrecht Institute. In short, cells were cultured with doxycycline for 6 weeks were resuspended and incubated with 2 ml of staining solution, consisting of 5 ug/ml Hoechst 34580 in cell culture medium. Cell pellet was resuspended in PBS. Cells were FACS sorted in 384-well plates and lysis of individual cells was performed for 2 h at 55°C using Proteinase K (Ambion) in 1x Cutsmart (New England Biolabs) followed by heat inactivation at 80°C for 10 min. The genomic DNA was subsequently fragmented with 200 nl 1 U NLAIII (New England Biolabs) in 1x Cutsmart (New England Biolabs) for 2 h at 37°C followed by heat inactivation at 65°C for 20 min. The genomic DNA was subsequently fragmented with 200 nl 1 U NLAIII (New England Biolabs) in 1x Cutsmart (New England Biolabs) for 2 h at 37°C followed by heat inactivation at 65°C for 20 min. Then, 50 nl of 50 mM barcoded double-stranded NLAIII adapters and 400 nl of 40 U T4 DNA ligase (New England Biolabs) in 1x T4 DNA ligase buffer (New England Biolabs) supplemented with 10 mM ATP (Invitrogen) was added to each well and ligated overnight at 16°C. Libraries were sequenced on an Illumina Nextseq 2000 with 1 × 50bp double-end sequencing. The fastq files were mapped to GRCH38 using the Burrows–Wheeler aligner. The mapped data were further analyzed using custom scripts in Python, which parsed for library barcodes, removed reads without a NlaIII sequence and removed PCR-duplicated reads. Copy number analysis was performed as described previously [58].

### RNA sequencing

For RNA sequencing, cells were cultured for 6 weeks with doxycycline. RNA was isolated using RNeasy Mini kit (Qiagen) and samples with low purity were further purified by isopropanol precipitation. RNA-seq libraries were prepared with the TruSeq Stranded Total RNA Library Prep Kit (Illumina) according to the manufacturer’s instructions. RNA-seq libraries were pooled and sequenced on a NextSeq2000 (Illumina) as 1 × 50 bp single end reads. RNA sequencing reads were aligned against human reference genome 37. The Bioconductor package DESeq2 was used to normalize raw read counts and to perform differential gene expression analysis [63].

### DNA methylation analysis

DNMT1 knockdown cells were cultured with doxycycline for 6 weeks. DNA was isolated using the DNeasy Blood & Tissue kit (Qiagen) followed by bisulfite conversion. The samples were subsequently run on an Infinium EPIC DNA methylation array. Data was analyzed with RNBeads [64].

### Nanopore sequencing of centromeres

For ONT sequencing of centromeres, pericentromeric regions were isolated by AlphaHOR-RES (alpha higher-order repeat restriction and enrichment by size)[65]. In short, genomic DNA was extracted from ∼25 million cells using an NEB High Molecular Weight DNA extraction kit followed by elution. Fully solubilized DNA was digested with MscI, and AseI. Digested DNA was loaded onto a 0.3% TAE agarose gel and run at 2 V/cm for 1 hour. Fragments larger than 20 kb were purified using a Zymoclean Large Fragment DNA Recovery Kit. DNA was subsequently prepared for DNA for sequencing using an ONT native library prep kit (LSK-109) and sequenced on a MinION (r9) flow cell. Nanopore raw data was basecalled using Guppy v5.0.17, and subsequently mapped against the CHM13 draft version 1.1 using minimap2 (map-ont settings). Reads were filtered for a mapping quality >=10. Methylation was then called using nanopolish 0.11.1 using the nanopolish call-methylation command. For binarized methylation calls, a cutoff of 2 was used so that scores >2 are interpreted as methylated and scores < -2 are interpreted as unmethylated.

### Sequencing and data analysis

For whole genome sequencing, DNA was isolated from cell pellets with the Qiasymphony (Qiagen) DNA isolation method and the Illumina TruSeq Nano DNA Library Prep Kit was used for library preparation. Samples were sequenced on HiSeq Xten or NovaSeq6000 platforms (Illumina) with 30x coverage. All samples were analysed with the HMF pipeline V4.8 (https://github.com/hartwigmedical/pipeline) which was locally deployed using GNU Guix with the recipe from https://github.com/UMCUGenetics/guix-additions. Full pipeline description is explained in [66], and details and settings of all the tools can be found at their Github page. Briefly, sequence reads were mapped against the human reference genome GRCh37 using Burrows-Wheeler Alignment (BWA-MEM) v0.7.5a [67]. Subsequently, somatic single base substitutions (SBSs), double base substitutions (DBSs) and small insertions and deletions (INDELS) were determined by Strelka v1.0.14 [68] that are further annotated by PURPLE. PURPLE (v2.53) combines B-allele frequency (BAF) from AMBER (v3.3), read depth ratios from COBALT (v1.7), and structural variants from GRIDSS[69] to estimate copy number profiles, variant allele frequency (VAF), variant clonality and microhomology context at the breakpoints. To obtain high-quality somatic mutations that can be attributed to in vivo mutagenesis in the ASC clones, we only considered somatic mutations with a PURPLE derived variant allele frequency higher than 30% as mutations that fall outside this range were potentially induced in vitro after the clonal passage. Analysis of the SVs was based on the LINX (v1.26)[70] output which interprets and annotates simple and complex SV events from PURPLE and GRIDSS output. LINX chains individual SVs into SV clusters and classifies these clusters into various event types. Clusters can have one SV (for simple events such as deletions and duplications which all have 1 clusterId), or multiple SVs, with ClusterId>1 and here considered as complex SV. We defined SV load as the total number of simple SV events. We quantified deletions and duplications (ResolvedType is ‘DEL’ or ‘DUP’) stratified by length (1–10 kb, 10–100 kb, 100kb–1Mb, 1–10Mb, >10 Mb). For complex SVs, we included “Complex_SV”, |”Complex_DEL”, “RECIP_INV” and “RECIP_TRANS” under resolved_Type annotation feature.

### Mutation burden analysis

The SBS, DBS and indel mutations were parsed from PURPLE vcfs by our developed R package Mutational Patterns[71] that was recently updated with DBS and indel functionality as well as COSMIC compatibility[72]. For each mutation type, we defined mutation burden as the total number of mutations of the autosomal genome.

## Author contributions

NB and JK performed wet-lab experiments. CV, AvH, and EK performed bioinformatic analyses. SB supported data management. EC and EK were involved in the conceptual design of the study. EC and JdR provided financial and lab support. EK wrote the paper. All authors proofread, made comments, and approved the paper.

## Acknowledgments

We are grateful for the support of Useq for facilitating the sequencing that was performed in this study and the Single Cell Sequencing Core facility at the Hubrecht Institute for their help with the single cell DNA sequencing. We would also like to thank Livio Kleij for his support with the microscopy and our colleagues from the genome diagnostics department for their support with the methylation arrays.

## References

1. Goldberg AD, Allis CD, Bernstein E. Epigenetics: a landscape takes shape. Cell. 2007;128: 635–638.

2. Mohn F, Schübeler D. Genetics and epigenetics: stability and plasticity during cellular differentiation. Trends Genet. 2009;25: 129–136.

3. Takahashi K, Tanabe K, Ohnuki M, Narita M, Ichisaka T, Tomoda K, et al. Induction of pluripotent stem cells from adult human fibroblasts by defined factors. Cell. 2007;131: 861–872.

4. Suvà ML, Riggi N, Bernstein BE. Epigenetic reprogramming in cancer. Science. 2013;339: 1567–1570.

5. Bergman Y, Cedar H. DNA methylation dynamics in health and disease. Nat Struct Mol Biol. 2013;20: 274–281.

6. Liu X, Gao Q, Li P, Zhao Q, Zhang J, Li J, et al. UHRF1 targets DNMT1 for DNA methylation through cooperative binding of hemi-methylated DNA and methylated H3K9. Nat Commun. 2013;4: 1563.

7. Brunetti L, Gundry MC, Goodell MA. DNMT3A in Leukemia. Cold Spring Harb Perspect Med. 2017;7. doi:10.1101/cshperspect.a030320

8. Cao T, Pan W, Sun X, Shen H. Increased expression of TET3 predicts unfavorable prognosis in patients with ovarian cancer-a bioinformatics integrative analysis. J Ovarian Res. 2019;12: 101.

9. Fang J-Y, Cheng Z-H, Chen Y-X, Lu R, Yang L, Zhu H-Y, et al. Expression of Dnmt1, demethylase, MeCP2 and methylation of tumor-related genes in human gastric cancer. World J Gastroenterol. 2004;10: 3394–3398.

10. Simó-Riudalbas L, Melo SA, Esteller M. DNMT3B gene amplification predicts resistance to DNA demethylating drugs. Genes Chromosomes Cancer. 2011;50: 527–534.

11. Langemeijer SMC, Kuiper RP, Berends M, Knops R, Aslanyan MG, Massop M, et al. Acquired mutations in TET2 are common in myelodysplastic syndromes. Nat Genet. 2009;41: 838–842.

12. Xiong Y, Dowdy SC, Xue A, Shujuan J, Eberhardt NL, Podratz KC, et al. Opposite alterations of DNA methyltransferase gene expression in endometrioid and serous endometrial cancers. Gynecol Oncol. 2005;96: 601–609.

13. Zhu Y-M, Huang Q, Lin J, Hu Y, Chen J, Lai M-D. Expression of human DNA methyltransferase 1 in colorectal cancer tissues and their corresponding distant normal tissues. Int J Colorectal Dis. 2007;22: 661–666.

14. Gaudet F, Hodgson JG, Eden A, Jackson-Grusby L, Dausman J, Gray JW, et al. Induction of tumors in mice by genomic hypomethylation. Science. 2003;300: 489–492.

15. Chen RZ, Pettersson U, Beard C, Jackson-Grusby L, Jaenisch R. DNA hypomethylation leads to elevated mutation rates. Nature. 1998;395: 89–93.

16. Karpf AR, Matsui S-I. Genetic disruption of cytosine DNA methyltransferase enzymes induces chromosomal instability in human cancer cells. Cancer Res. 2005;65: 8635–8639.

17. Sheaffer KL, Elliott EN, Kaestner KH. DNA Hypomethylation Contributes to Genomic Instability and Intestinal Cancer Initiation. Cancer Prev Res. 2016;9: 534–546.

18. Chen T, Hevi S, Gay F, Tsujimoto N, He T, Zhang B, et al. Complete inactivation of DNMT1 leads to mitotic catastrophe in human cancer cells. Nat Genet. 2007;39: 391–396.

19. Barra V, Schillaci T, Lentini L, Costa G, Di Leonardo A. Bypass of cell cycle arrest induced by transient DNMT1 post-transcriptional silencing triggers aneuploidy in human cells. Cell Div. 2012;7: 2.

20. Costa G, Barra V, Lentini L, Cilluffo D, Di Leonardo A. DNA demethylation caused by 5-Aza-2’-deoxycytidine induces mitotic alterations and aneuploidy. Oncotarget. 2016;7: 3726–3739.

21. Saksouk N, Simboeck E, Déjardin J. Constitutive heterochromatin formation and transcription in mammals. Epigenetics & Chromatin. 2015. doi:10.1186/1756-8935-8-3

22. Scelfo A, Fachinetti D. Keeping the Centromere under Control: A Promising Role for DNA Methylation. Cells. 2019;8. doi:10.3390/cells8080912

23. Berman BP, Weisenberger DJ, Aman JF, Hinoue T, Ramjan Z, Liu Y, et al. Regions of focal DNA hypermethylation and long-range hypomethylation in colorectal cancer coincide with nuclear lamina-associated domains. Nat Genet. 2011;44: 40–46.

24. Timp W, Bravo HC, McDonald OG, Goggins M, Umbricht C, Zeiger M, et al. Large hypomethylated blocks as a universal defining epigenetic alteration in human solid tumors. Genome Med. 2014;6: 61.

25. Jafri MA, Ansari SA, Alqahtani MH, Shay JW. Roles of telomeres and telomerase in cancer, and advances in telomerase-targeted therapies. Genome Med. 2016;8: 69.

26. Rodriguez-Martin B, Alvarez EG, Baez-Ortega A, Zamora J, Supek F, Demeulemeester J, et al. Pan-cancer analysis of whole genomes identifies driver rearrangements promoted by LINE-1 retrotransposition. Nat Genet. 2020;52: 306–319.

27. Saha AK, Mourad M, Kaplan MH, Chefetz I, Malek SN, Buckanovich R, et al. The Genomic Landscape of Centromeres in Cancers. Sci Rep. 2019;9: 11259.

28. Slee RB, Steiner CM, Herbert B-S, Vance GH, Hickey RJ, Schwarz T, et al. Cancer-associated alteration of pericentromeric heterochromatin may contribute to chromosome instability. Oncogene. 2012;31: 3244–3253.

29. Tate JG, Bamford S, Jubb HC, Sondka Z, Beare DM, Bindal N, et al. COSMIC: the Catalogue Of Somatic Mutations In Cancer. Nucleic Acids Res. 2019;47: D941–D947.

30. Degasperi A, Zou X, Amarante TD, Martinez-Martinez A, Koh GCC, Dias JML, et al. Substitution mutational signatures in whole-genome-sequenced cancers in the UK population. Science. 2022;376. doi:10.1126/science.abl9283

31. Alexandrov LB, Kim J, Haradhvala NJ, Huang MN, Tian Ng AW, Wu Y, et al. The repertoire of mutational signatures in human cancer. Nature. 2020;578: 94–101.

32. Li Y, Roberts ND, Wala JA, Shapira O, Schumacher SE, Kumar K, et al. Patterns of somatic structural variation in human cancer genomes. Nature. 2020;578: 112–121.

33. Chen W-H, Lu G, Chen X, Zhao X-M, Bork P. OGEE v2: an update of the online gene essentiality database with special focus on differentially essential genes in human cancer cell lines. Nucleic Acids Res. 2017;45: D940–D944.

34. Hart T, Chandrashekhar M, Aregger M, Steinhart Z, Brown KR, MacLeod G, et al. High-Resolution CRISPR Screens Reveal Fitness Genes and Genotype-Specific Cancer Liabilities. Cell. 2015;163: 1515–1526.

35. Horlbeck MA, Gilbert LA, Villalta JE, Adamson B, Pak RA, Chen Y, et al. Compact and highly active next-generation libraries for CRISPR-mediated gene repression and activation. Elife. 2016;5. doi:10.7554/eLife.19760

36. Ho S-M, Hartley BJ, Flaherty E, Rajarajan P, Abdelaal R, Obiorah I, et al. Evaluating Synthetic Activation and Repression of Neuropsychiatric-Related Genes in hiPSC-Derived NPCs, Neurons, and Astrocytes. Stem Cell Reports. 2017;9: 615–628.

37. Kaelin WG Jr. Molecular biology. Use and abuse of RNAi to study mammalian gene function. Science. 2012;337: 421–422.

38. Maslov AY, Lee M, Gundry M, Gravina S, Strogonova N, Tazearslan C, et al. 5-aza-2’-deoxycytidine-induced genome rearrangements are mediated by DNMT1. Oncogene. 2012;31: 5172–5179.

39. Santi DV, Norment A, Garrett CE. Covalent bond formation between a DNA-cytosine methyltransferase and DNA containing 5-azacytosine. Proc Natl Acad Sci U S A. 1984;81: 6993–6997.

40. Kuijk E, Jager M, van der Roest B, Locati MD, Van Hoeck A, Korzelius J, et al. The mutational impact of culturing human pluripotent and adult stem cells. Nat Commun. 2020;11: 2493.

41. Kuijk E, Blokzijl F, Jager M, Besselink N, Boymans S, Chuva de Sousa Lopes Sm, et al. Early divergence of mutational processes in human fetal tissues. Sci Adv. 2019;5: eaaw1271.

42. Jager M, Blokzijl F, Kuijk E, Bertl J, Vougioukalaki M, Janssen R, et al. Deficiency of nucleotide excision repair is associated with mutational signature observed in cancer. Genome Res. 2019;29: 1067–1077.

43. Christensen S, Van der Roest B, Besselink N, Janssen R, Boymans S, Martens JWM, et al. 5-Fluorouracil treatment induces characteristic T>G mutations in human cancer. Nat Commun. 2019;10: 4571.

44. Blokzijl F, de Ligt J, Jager M, Sasselli V, Roerink S, Sasaki N, et al. Tissue-specific mutation accumulation in human adult stem cells during life. Nature. 2016;538: 260–264.

45. Nguyen L, Jager M, Lieshout R, de Ruiter PE, Locati MD, Besselink N, et al. Precancerous liver diseases do not cause increased mutagenesis in liver stem cells. Commun Biol. 2021;4: 1301.

46. Wang K-Y, James Shen C-K. DNA methyltransferase Dnmt1 and mismatch repair. Oncogene. 2004;23: 7898–7902.

47. Drost J, van Boxtel R, Blokzijl F, Mizutani T, Sasaki N, Sasselli V, et al. Use of CRISPR-modified human stem cell organoids to study the origin of mutational signatures in cancer. Science. 2017. pp. 234–238. doi:10.1126/science.aao3130

48. Zhang C-Z, Spektor A, Cornils H, Francis JM, Jackson EK, Liu S, et al. Chromothripsis from DNA damage in micronuclei. Nature. 2015;522: 179–184.

49. Eden A, Gaudet F, Waghmare A, Jaenisch R. Chromosomal instability and tumors promoted by DNA hypomethylation. Science. 2003;300: 455.

50. Nurk S, Koren S, Rhie A, Rautiainen M, Bzikadze AV, Mikheenko A, et al. The complete sequence of a human genome. Science. 2022;376: 44–53.

51. Pappalardo XG, Barra V. Losing DNA methylation at repetitive elements and breaking bad. Epigenetics Chromatin. 2021;14: 25.

52. Wong N, Lam WC, Lai PB, Pang E, Lau WY, Johnson PJ. Hypomethylation of chromosome 1 heterochromatin DNA correlates with q-arm copy gain in human hepatocellular carcinoma. Am J Pathol. 2001;159: 465–471.

53. Qu GZ, Grundy PE, Narayan A, Ehrlich M. Frequent hypomethylation in Wilms tumors of pericentromeric DNA in chromosomes 1 and 16. Cancer Genet Cytogenet. 1999;109: 34–39.

54. Feng S, Jacobsen SE, Reik W. Epigenetic Reprogramming in Plant and Animal Development. Science. 2010. pp. 622–627. doi:10.1126/science.1190614

55. Jönsson ME, Ludvik Brattås P, Gustafsson C, Petri R, Yudovich D, Pircs K, et al. Activation of neuronal genes via LINE-1 elements upon global DNA demethylation in human neural progenitors. Nat Commun. 2019;10: 3182.

56. García-Muse T, Aguilera A. R Loops: From Physiological to Pathological Roles. Cell. 2019. pp. 604–618. doi:10.1016/j.cell.2019.08.055

57. Gonzalo S, Jaco I, Fraga MF, Chen T, Li E, Esteller M, et al. DNA methyltransferases control telomere length and telomere recombination in mammalian cells. Nat Cell Biol. 2006;8: 416–424.

58. Bakker B, Taudt A, Belderbos ME, Porubsky D, Spierings DCJ, de Jong TV, et al. Single-cell sequencing reveals karyotype heterogeneity in murine and human malignancies. Genome Biol. 2016;17: 115.

59. Sanjana NE, Shalem O, Zhang F. Improved vectors and genome-wide libraries for CRISPR screening. Nat Methods. 2014;11: 783–784.

60. Ran FA, Ann Ran F, Hsu PD, Wright J, Agarwala V, Scott DA, et al. Genome engineering using the CRISPR-Cas9 system. Nature Protocols. 2013. pp. 2281–2308. doi:10.1038/nprot.2013.143

61. Conant D, Hsiau T, Rossi N, Oki J, Maures T, Waite K, et al. Inference of CRISPR Edits from Sanger Trace Data. CRISPR J. 2022. doi:10.1089/crispr.2021.0113

62. Livak KJ, Schmittgen TD. Analysis of relative gene expression data using real-time quantitative PCR and the 2(-Delta Delta C(T)) Method. Methods. 2001;25: 402–408.

63. Eccles DA. Creating Differential Transcript Expression Results with DESeq2 v2. doi:10.17504/protocols.io.8epv51686l1b/v2

64. Müller F, Scherer M, Assenov Y, Lutsik P, Walter J, Lengauer T, et al. RnBeads 2.0: comprehensive analysis of DNA methylation data. Genome Biology. 2019. doi:10.1186/s13059-019-1664-9

65. Altemose N, Maslan A, Smith O, Sundararajan K, Brown R, Straight A, et al. AlphaHOR-RES: a method for enriching centromeric DNA v1. protocols.io. ZappyLab, Inc.; 2021. doi:10.17504/protocols.io.bv9vn966

66. Priestley P, Baber J, Lolkema M, Steeghs N, de Bruijn E, Duyvesteyn K, et al. Pan-cancer whole genome analyses of metastatic solid tumors. 2018. doi:10.1101/415133

67. Li H, Durbin R. Fast and accurate short read alignment with Burrows-Wheeler transform. Bioinformatics. 2009;25: 1754–1760.

68. Saunders CT, Wong WSW, Swamy S, Becq J, Murray LJ, Cheetham RK. Strelka: accurate somatic small-variant calling from sequenced tumor-normal sample pairs. Bioinformatics. 2012;28: 1811–1817.

69. Cameron DL, Baber J, Shale C, Valle-Inclan JE, Besselink N, van Hoeck A, et al. GRIDSS2: comprehensive characterisation of somatic structural variation using single breakend variants and structural variant phasing. bioRxiv. bioRxiv; 2020. doi:10.1101/2020.07.09.196527

70. Cameron DL, Baber J, Shale C, Papenfuss AT, Valle-Inclan JE, Besselink N, et al. GRIDSS, PURPLE, LINX: Unscrambling the tumor genome via integrated analysis of structural variation and copy number. bioRxiv. 2019. p. 781013. doi:10.1101/781013

71. Blokzijl F, Janssen R, van Boxtel R, Cuppen E. MutationalPatterns: comprehensive genome-wide analysis of mutational processes. Genome Med. 2018;10: 33.

72. Manders F, Brandsma AM, de Kanter J, Verheul M, Oka R, van Roosmalen MJ, et al. MutationalPatterns: The one stop shop for the analysis of mutational processes. bioRxiv. 2021. p. 2021.11.01.466730. doi:10.1101/2021.11.01.466730

73. Ge SX, Jung D, Yao R. ShinyGO: a graphical gene-set enrichment tool for animals and plants. Bioinformatics. 2019;36: 2628–2629.

